# Bedtk: Finding Interval Overlap with Implicit Interval Tree

**DOI:** 10.1101/2020.07.07.190744

**Authors:** Heng Li, Jiazhen Rong

## Abstract

**Summary:** We present bedtk, a new toolkit for manipulating genomic intervals in the BED format. It supports sorting, merging, intersection, subtraction and the calculation of the breadth of coverage. Bedtk employs implicit interval tree, a new data structure for fast interval overlap queries. It is several to tens of times faster than existing tools and tends to use less memory.

**Availability:** https://github.com/lh3/bedtk

## 1 Introduction

Processing genomic intervals is a routine task and has many practical applications in bioinformatics. BEDTools (Quinlan and Hall, 2010) is a popular toolkit to manipulate intervals in the BED format. However, it can be slow given large datasets. BEDOPS (Neph et al., 2012) addresses this issue by streaming sorted BED files. While this strategy improves performance, it is less convenient to use and is limited to a subset of interval operations. These observations motivated us to develop bedtk that achieves high performance without requiring sorting.

## 2 Methods

Efficiently finding overlapping intervals is a core functionality behind all interval processing tools. Bedtk introduces implicit interval tree, a novel data structure to perform interval overlap queries against a static list of intervals.

### 2.1 Implicit binary search trees

A binary search tree (BST) is a binary tree where each node is associated with a key and this key is no less than all keys in the left subtree and no greater than all keys in the right subtree. Here we show that a BST can be implicitly represented by a sorted array. In this implicit BST, each node is an element in the array and the tree topology is inferred from the index of each element.

More specifically, consider a sorted array consisting of 2^*k*+1^ − 1 elements. This array can implicitly represent a binary tree with *K* + 1 levels, with leaves put at level 0 and the root at level *K* (Fig. 1). This implicit BST has the following properties:

1. At level *k*, the first node is indexed by 2^*k*^ − 1. As a special case, the root of the tree is indexed by 2^*K*^ − 1.
2. For a node indexed by *x* at level *k*, its left child is indexed by *x* − 2^*k*−1^, and its right child indexed by *x* + 2^*k*−1^.
3. For a node indexed by *x* at level *k*, it is a left child if its (*k* + 1)-th bit is 0 (i.e. ⌊*x*/2^*k*+1^⌋ is an even number), and its parent node is *x* + 2^*k*^. Node *x* is a right child if its (*k* + 1)-th bit is 1 (i.e. ⌊*x*/2^*k*+1^⌋ is an odd number), and its parent node is *x* − 2^*k*^.
4. For a node indexed by *x* at level *k*, there are 2^*k*+1^ − 1 nodes descending from *x* (including *x*). The left-most leaf in this subtree is *x* − (2^*k*^ − 1).

**Fig. 1.**
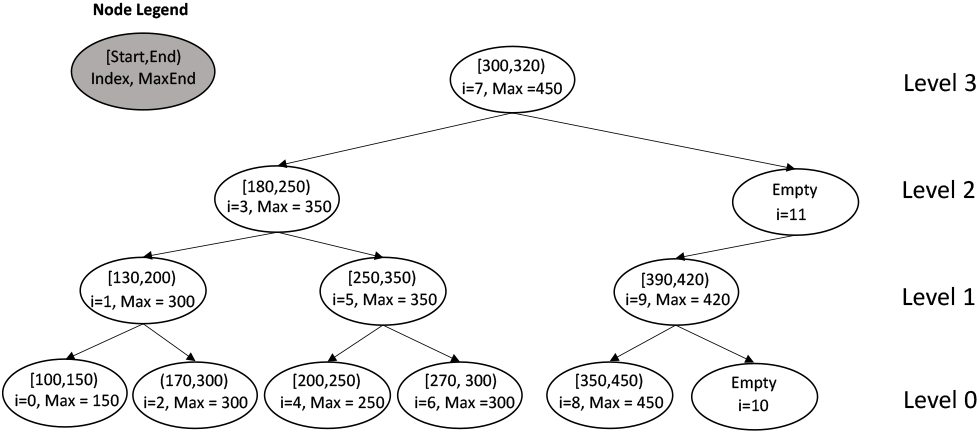
Example of implicit interval tree.

With these properties, we can identify the parent and the children of a node in *O*(1) time. Fig. 1 shows an example of implicit BST for the start positions of a list of intervals which is sorted by the start positions. When there are fewer than 2^*k*+1^ − 1 elements in the array, we still assume a full binary tree with some nodes set to “empty”. Notably in this case, an empty node may have a non-empty child (e.g. in Fig. 1, empty node 11 has a non-empty child node 9).

### 2.2 Implicit interval trees

An augmented interval tree is a data structure for interval overlap queries. It is an extension to BST where a node corresponds to an interval and each node additionally keeps a MaxEnd field which is the largest end position in the subtree descending from the node. Because a BST can be implicitly represented with an array, an augmented interval tree can be represented with an array as well (Fig. 1).

## 3 Results

We implemented implicit interval tree in bedtk along with few other common operations such as sorting and interval merging. We compared bedtk to BEDTools and BEDOPS on two BED files: 1) 1,194,285 exons from GenCode annotations and 2) 9,944,559 alignment intervals of long RNA-seq reads. Both are available at https://github.com/lh3/cgranges/releases.

As is shown in Table 1, bedtk is consistently faster than BEDTools and BEDOPS for all the evaluated operations, even when we discount sorting time for the BEDOPS “intersection” operation. The performance gap between bedtk and BEDTools is even larger for unsorted input (“intersect” and “coverage”). This exemplifies the advantage of the implicit interval tree. BEDOPS takes the least memory for sorting potentially because it uses advanced encoding; it uses even less memory for “intersect” as it streams the input instead of loading one or both input files into memory. However, requiring sorted input complicates data processing pipelines and wastes working disk space. And even with sorted input, BEDOPS is slower than bedtk.

**Table 1.** Performance of interval operations

| Operation | bedtk | BEDTools | BEDOPS |
| --- | --- | --- | --- |
| sort | 5.35s / 393MB | 21.14s / 2731MB | 11.25s / 205MB |
| intersect | 6.42s / 19MB | 267.32s / 397MB | 8.78s / 3MB |
| coverage | 10.91s / 19MB | 257.11s / 678MB |  |

Each cell gives the CPU time in sections and peak memory in megabytes (MB). The “sort” operation sorts file 2. “intersect” reports intervals in file 2 that overlaps intervals in file 1. For each interval in file 2, “coverage” computes the number of bases covered by intervals in file 1. BEDOPS does not support the “coverage” operation. It also requires sorted input for “intersect”; sorting time is excluded.

## 4 Discussions

An implicit interval tree can be implemented in <100 lines of C++ code. It is a simple yet efficient data structure for fast interval queries. Daniel Jones alters the memory layout of implicit interval tree for improved cache locality (https://github.com/dcjones/coitrees). It speeds up query at the cost of more memory and unsorted query output. Meanwhile, Mike Lin gives up the standard top-down interval tree query and instead allows to start a query from any node in the tree (https://github.com/mlin/iitii). This improves the performance on practical data. In additional to interval trees, nested containment list (Alekseyenko and Lee, 2007) and augmented interval list (Feng et al, 2019) are alternative data structures for fast interval overlap queries. However, no standalone user-oriented tools have implemented these advanced algorithms. Bedtk is the first toolkit that is designed for end users and outperforms popular tools for common interval operations.

## Acknowledgements

We thank Daniel C. Jones and Mike Lin for further improving the performance of implicit interval trees.

## References

Alekseyenko, A.V. and Lee, C.J. (2007). Nested Containment List (NCList): a new algorithm for accelerating interval query of genome alignment and interval databases, Bioinformatics, 23:1386–1393

Feng J., Ratan A. and Sheffield N.C. (2019). Augmented Interval List: a novel data structure for efficient genomic interval search, Bioinformatics, 35:4907–4911

Neph S. et al. (2012). BEDOPS: high-performance genomic feature operations, Bioinformatics, 28: 1919–1920

Quinlan A.R. and Hall I.M. (2010). BEDTools: a flexible suite of utilities for comparing genomic features. Bioinformatics, 26: 841–842

